# Pi16^+^ fibroblast-derived Csf1 shapes skin topography

**DOI:** 10.64898/2026.04.02.716114

**Authors:** Anthony Altieri, Erika E. McCartney, Shaheed W. Hakim, Jean X. Jiang, Matthew B. Buechler

## Abstract

*Peptidase inhibitor 16* (*Pi16*)-expressing fibroblasts are found across tissues and species, but their functional role is unclear. As fibroblasts and macrophages have been proposed to exist in a reciprocal circuit, we hypothesized *Pi16*^+^ fibroblasts may regulate macrophage homeostasis. Flow cytometry revealed ∼80% of skin fibroblasts express *Pi16*, leading us to investigate the role of these cells in maintaining a macrophage niche in this tissue. We generated an *in vivo* system where fibroblast-derived *Colony Stimulating Factor 1* (*Csf1)* was constitutively eliminated in *Pi16*^*+*^ fibroblasts by crossing animals with a *Csf1fl/fl* allele to mice in which the gene *Pi16* drives an IresCre cassette. Deletion of *Csf1* in *Pi16*^+^ fibroblasts resulted in significant diminishment of CD64^+^ and CD11c^+^ macrophages alongside expansion of PDPN^+^YFP^+^ fibroblasts. Alterations in cell population dynamics coincided with thickening of both the dermis and fascial compartments of the skin. Deletion of *Csf1* in *Pi16*^+^ fibroblasts delayed early wound healing in a unsplinted mouse model. Loss of *PI16*^+^ fibroblasts was observed in individuals with limited (lSSc) and diffuse (dSSc) systemic Scleroderma compared to healthy controls. These findings suggest that loss of *Csf1* in *Pi16*^+^ fibroblasts elicit changes in the population dynamics of skin macrophages and modifications to tissue architecture.

## Introduction

The skin is the largest organ in the human body and acts as the vanguard between internal environments and external threats by orchestrating interactions between stromal and immune cells^1,2^. Fibroblasts are stromal cells which regulate local tissue architecture and resident immune cell populations by producing extracellular matrix (ECM) and cytokines^3^, including the macrophage trophic factor *Colony Stimulating Factor 1* (*Csf1*)^4,5^. Macrophages are heterogeneous immune cells of the myeloid lineage which express *CSF1 receptor* (*Csf1r)* and orchestrate both inflammatory and tissue repair responses in response to tissue damage through a variety of means including producing cytokines that can alter fibroblast responses, such as Transforming Growth Factor (TGF)-β, platelet-derived growth factors (PDGF)-α and -β, and insulin-like growth factor (IGF)-1^6–9^. As a result, fibroblasts and macrophages have been posited to exist in a bidirectional two-cell circuit anchored by the *Csf1*-*Csf1r* axis across tissues^10^. The fibroblast:macrophage two-cell circuit concept has been studied in silico^11^, in vitro^12^ and in vivo leveraging stroma-targeted in vivo cre-loxP systems^7,13–16^ in several tissue contexts.

In the skin microenvironment, conditional deletion of *Csf1* in *Col1a2*+ mesenchymal cells resulted in CCR2^+^ and CCR2^-^ macrophage abrogation in the adventitia^7^ (hereafter referred to as fascia). *Dermatopontin* (*Dpt)*-expressing (*Dpt*)^+^ fibroblasts are found across human and mouse tissues, are composed of two substate fibroblast populations defined by the expression of *Peptidase inhibitor 16* (*Pi16)* and *Collagen 15a1* (*Col15a1)* and express *Csf1*^17^. We recently utilized the *Dpt*^IresCreERT2^ genetic tool^17^ to reveal that *Dpt*^+^ fibroblasts control macrophage homeostasis, tissue integrity and responses to wounding in the skin^18^. Additionally, our work revealed that alterations to the skin niche upon loss of *Csf1* in *Dpt*^+^ fibroblasts caused transcriptional and numeric changes in skin fibroblasts, including possible alterations to *Pi16+* fibroblasts^18^. Finally, dysregulation of the *Dpt*^+^ fibroblast:macrophage axis was observed in the skin of patients with systemic scleroderma (SSc) compared to healthy controls^18^.

The precise contribution of the *Pi16*^+^ fibroblast substate in reciprocal regulation of the fibroblast:macrophage *Csf1*-dependent cell-circuit has not been directly tested. We investigated the *Pi16*^+^ fibroblast:macrophage circuit in the skin using the *Pi16*^IresCre^ genetic model. We hypothesized that *Csf1*-driven *Pi16*^+^ fibroblast:macrophage interactions are critical for skin homeostasis. We pursued this question through conditional deletion of *Csf1* expression in *Pi16*^+^ fibroblasts in vivo and performing flow cytometric, histological, and immunofluorescence analyses. Here, we demonstrate that *Pi16*^+^ fibroblasts maintain an essential macrophage niche in the skin through *Csf1* provision. Diminishment of macrophages in response to *Csf1* loss was coincident with increased *Pi16*^*+*^ fibroblast abundance in the fascia of the skin, increased fascial thickness, and delays in early wound healing kinetics in the skin. Perturbation of the *PI16*^+^ fibroblast:macrophage axis occurred in human disease, where loss of *PI16*^+^ fibroblasts and was observed in the skin of patients with SSc compared with healthy controls.

## Methods

### Mice and in vivo treatments

*Pi16*^IresCre^ *(Pi16)* mice were designed, generated and bred at the Centre for Phenogenomics (TCP). *Csf1(exons 4-6)*^*fl/fl*^ mice with exons 4-6 of the *Csf1* gene flanked by *loxP* sites were donated by S. Werner (University of Texas Health Center, San Antonio)^19^ via C. Robbins (University of Toronto) and crossed with *Pi16*^IresCre^ at TCP, resulting in Csf1-knockout mice *(cre*^*+/-*^; *Csf1*^*fl/fl*^ or *cre*^*+/+*^; *Csf1*^*fl/fl*^). For mouse experiments, littermate *cre*^*+/*^ or *cre*^*+/+*^; *Csf1*^*fl/+*^ or *Csf1*^*fl/fl*^ were used as controls. Male and female mice between 8-12 weeks of age were used for all studies. Mice were housed in specific-pathogen free, individually ventilated cages. Animal rooms were maintained on a 13:11 hour light-dark cycle and were temperature controls at ∼22°C. All mouse studies were approved by the Animal Care Committee at the TCP and the University of Toronto. No statistical methods were used to predetermine sample size. Experiments were not randomized and investigators were not blinded for allocation of experiments and outcome assessments except for histological analyses and wound healing assays, where investigators were blinded to the background of mice for manual annotation of tissue thickness and wound boundaries.

### Mouse tissue digestions and stromal cell isolation

Flank skin tissue was processed as previously described^18^ by shaving hair, removing adipose tissue, and mincing tissue in RPMI + 2% FBS with dispase (800 ul ml^-1^; Life Technologies, 1010459001), collagenase P (200 ul ml^-1^; Roche, 1129002001), and DNAse1 (100 ul ml^-1^; Roche 10104159001). Digestion at 37°C (water bath) involved three 15-minute incubations with fractions filtered (70 μm) after each incubation period. Cells were centrifuged (1600 RPM, 2 minutes), blocked with Fc Block (Biolegend, 101302 (1:1000)) and labeled with following antibodies (dilution; catalog #; supplier in parenthesis): BUV395 CD45 (1:300; 564279; BD Biosciences); BUV737 CD11b (1:400; 612800; BD Biosciences); BV421 CD64 (1:800; 139309; Biolegend); BV605 GR1 (1:200; 108439; Biolegend); PE-Cy7 Gr1 (1:400; 108416; Biolegend); BV650 F4/80 (1:100; 123149; Biolegend); BV711 CD206 (1:800; 141727; Biolegend); APC CCR2 (1:10; FAB5538a; R&D systems); PE-Cy7 CD11c (1:100; 117318; Biolegend); PE-Cy7 TIM4 (1:200; 130009; Biolegend); BV421 LY6C (1:200; 128031; Biolegend); APC F4/80 (1:800; 157306; Biolegend); BUV395 CD9 (1:100; 740244; BD Biosciences); BUV737 CD90.2 (1:800; 741702; BD Biosciences); SB600 PDPN (1:200; 63538182; eBioscience); BV711 SCA1 (1:400; 108131; Biolegend); BV786 CD54 (1:400; 740855; BD Biosciences); PCP LY6C (1:800; 128012; Biolegend); PE-Cy7 CD31 (1:200; 102524; Biolegend); PE-Cy7 CD45 (1:400; 103114; Biolegend); PE-Cy7 EpCAM (1:400; 118216;Biolegend); APC CD26 (1:200;137807; Biolegend); APC-Cy7 CD201 (1:200;). Dead cells were excluded with the Zombie Violet fixable viability dye (1:1000; 77477; Biolegend) or 7-AAD Viability Solution (1:400; 420404; Biolegend). Data were acquired using the BD FACSymphony A5 SE and analyzed using FlowJo v10.9.0.

### Tissue processing for cDNA synthesis and quantitative reverse transcriptase (qRT)-PCR

Mouse flank skin was isolated by shaving hair and removing adipose tissue. Flank skin samples were homogenized in 2.8 mm Ceramic PowerBead Tubes (Qiagen) containing 350 uL RNA lysis solution (1:100 Buffer RLT (Qiagen):2-Mercaptoethanol (Sigma M3148)) at 2.9 m/s for 30 seconds using a Bead Ruptor Elite (Omi International). RNA was extracted from cell lysates using after centrifugation at 8000xg for 3 mins using a RNAeasy Mini Kit (Qiagen). The quantity and quality of Total RNA was measured using the NanoDrop Eight Spectrophotometer (ThermoFisher Scientific). 100 ng total RNA was used for cDNA synthesis. Reverse transcription was performed with 200 ng of RNA ABI advanced kit including RNase inhibitor (ABI Cat. 4374966). qPCR was performed on cDNA product diluted with RNase-free water (1:100) using Taqman Gene Expression Assay probe for the *Csf1 (Mm00432686). Gapdh* (Mm99999915_g1) was used as the reference gene. Taqman Fast Advanced Master Mix (Thermofisher) was used to dilute each probe (1:10) and qPCR was performed with the following cycling parameters: 50°C for 2 min, 95°C for 20 sec, and then repeating 40 cycles, 95°C for 1 sec, 60°C for 20 sec. All qPCR reactions were conducted in duplicate in a MicroAmp optical 96-well reaction plate using QuantStudio 6Pro (Applied Biosystems). The average of threshold cycle (Ct) of target gene(s) was normalized to *Gapdh*.

### Full thickness, unsplinted wounding

Mice were anesthetized using isoflurane and the dorsal portion of the mouse back was shaved from the scapular area to the lumbar area. Residual hair was removed using Nair and rinsed with sterile alcohol pads containing 2-propanol 70% v/v (Alliance). On the day of wounding mice were first anesthetized using isoflurane in an induction chamber then moved to a nose cone placed on a heated pad to enable wound generation under anesthesia. A 6 mm punch biopsy (VWR) was used to generate the outline of the wound and skin tissue excision was performed using surgical scissors. Wounds were dressed in Tegaderm (3M) and changed every day. Wound images were acquired using a digital camera on a fixed camera stand. Wound images were manually traced, calculated, and analyzed for percent closure using Image J/FIJI.

### Tissue processing and histology

Skin tissue samples were formalin fixed, paraffin-embedded and sectioned (5 μM) and sections were stained with Hematoxylin and Eosin. Slides were scanned at 20X using an Olympus VS120 slide scanner equipped with a Hamamatsu ORCA-R2 C10600 digital camera (Evident).

### Quantification of Immunofluorescence

Mouse skin samples in formalin were submitted to the TCP Pathology core for histology and immunohistochemistry. The samples were processed in a TissueTek VIP 6 tissue processor (Sakura Finetek), paraffin-embedded and 5 µm-thick sections were collected onto charged slides (Assure, Epic Scientific). The sections were deparaffinized in a TissueTek Prisma instrument (Sakura Finetek), submitted to heat-induced epitope retrieval with citrate buffer pH 6.0 (10mM Sodium Citrate, 0.05% Tween-20) and incubated in a pressure cooker for 10 minutes, followed for 30 minutes at room temperature. The sections were incubated in Protein Block (Agilent) to minimize non-specific antibody binding before each primary antibody incubation. Primary antibodies goat anti-GFP (1:500, Abcam) and rat anti-F4/80 (1:100, Abcam) were diluted in Antibody Diluent solution (Agilent) and incubated sequentially. Slides were incubated overnight in anti-GFP primary antibody at 4°C. On the following day, the tissue sections were washed in 1x TRIS-buffered saline with 0.1% Tween-20 (TBST), then incubated in anti-goat IgG conjugated with Alexa Fluor 647 (Invitrogen) diluted at 1:200 in Antibody Diluent for 1 hour at room temperature. After rinsing in TBST, the sections were incubated in Protein Block prior to overnight incubation in anti-F4/80 primary antibody overnight at 4°C. On the following day, tissue sections were washed in TBST, incubated in anti-rat IgG conjugated with Alexa Fluor 555 (Invitrogen) diluted at 1:200 in Antibody Diluent for 1 hour at room temperature, rinsed in TBST then counterstained with DAPI (Sigma). Tissue autofluorescence was quenched in Sudan black B saturated solution in 70% ethanol (Sigma) for 25 minutes at room temperature, then coverslips were mounted with Vectashield Vibrance anti-fade mounting medium (Vector, H-1700). Slides were scanned at 20X using an Olympus VS120 slide scanner equipped with a Hamamatsu ORCA-R2 C10600 digital camera (Evident). Image analysis was performed using HALO software (Indica Labs). Cell types were identified using the HighPlex-FL algorithm and fine-tuned for cell types of interest detailed in the main texts. Regions of interest, including dermis, panniculus muscle (PM), dermal white adipose tissue, and fascia were manually annotated based on histological appearance of skin tissue sections.

### Analysis of human systemic scleroderma single-cell and bulk RNA sequencing datasets

Human single cell sequencing data and metadata^20^ was downloaded from Gene Expression Omnibus (GEO) with ascension number GSE195452, using metadata containing additional cell information such as patient diagnosis, including healthy, lSSc, or dSSs. Subsequent data analyses were performed in R Studio (v.4.4.1) using Seurat (v.5.3.0). Unannotated cells (denoted in metadata) with more that 25% mitochondrial genes were removed. Gene expression of the remaining cells was log normalized and the top 3000 highly variable genes were detected using FindVariableFeatures(). Principal component analysis was used for dimensionality reduction. Nearest neighbors were found, and cells were clustered using the first 30 principal components and a resolution of 0.5, respectively. Differential abundance (DA) analysis was performed using the R package MiloR (v.2.5.1), by which significant neighborhoods were identified using spatial FDR correction. Data were visualized by overlaying DA results onto previously generated UMAPs.

### Quantification, statistical analysis, and data presentation

Data analysis was performed as indicated in the figure legends. Statistical analysis and data presentation were performed using GraphPad Prism 10.6.1. Exact p-values are reported, otherwise significance is indicated as *p ≤ 0.05, **p ≤ 0.01, ***p ≤ 0.001, ****p ≤ 0.001.

## Results

### Pi16^+^ fibroblasts represent 80% of skin fibroblasts and are a critical source of skin Csf1

*Pi16*^+^ fibroblasts are a transcriptional subset of *Dpt*^+^ fibroblasts^17^, which act as a non-redundant source of *Csf1* in steady-state skin^18^. To directly investigate whether CSF1 production by *Pi16*^+^ fibroblasts maintains macrophage homeostasis, we generated an *in vivo* system where fibroblast-derived *Csf1* was constitutively eliminated by crossing the *Csf1fl/fl Exons* 5–7 allele^18,19^ to animals in which the gene *Pi16* drives an IresCre cassette *Pi16*^IresCre^ (*Pi16*^IresCre^;*Rosa26*^lox.stop.lox-YellowFluorescentProtein^*Csf1*^FLOX^; *Pi16*^IresCre^;*R26*^YFP^;*Csf1*^fl/fl^ (**Fig. 1A**). We observed robust YFP^+^ expression in the skin where flow cytometric analysis demonstrated that *Pi16*^+^ fibroblasts represent ∼80% of all skin fibroblasts (**Fig. 1B-C)**, as enumerated as non-hematopoetic (CD45), - epithelial (EpCAM), -endothelial (CD31) cells that express podoplanin (PDPN). Subsequent characterization of YFP^+^ skin fibroblasts identified higher abundance of CD26, CD90, LY6C, and SCA1 markers compared to YFP^-^ fibroblasts, suggesting fibroblasts marked by YFP in this system exhibit a distinct surface phenotype (**Fig. 1D-E**). qPCR analysis of whole skin tissue confirmed a statistically significant reduction of *Csf1* (2.3-fold) in *Pi16*^IresCre^;*R26*^YFP^;*Csf1*^fl/fl^ compared to *Pi16*^IresCre^;*R26*^YFP^;*Csf1*^CTRL^ animals (**Fig. 1F)**, confirming the efficacy of the model and suggesting that *Pi16*^+^ fibroblasts are an essential source of *Csf1* in the skin.

**Figure 1.**
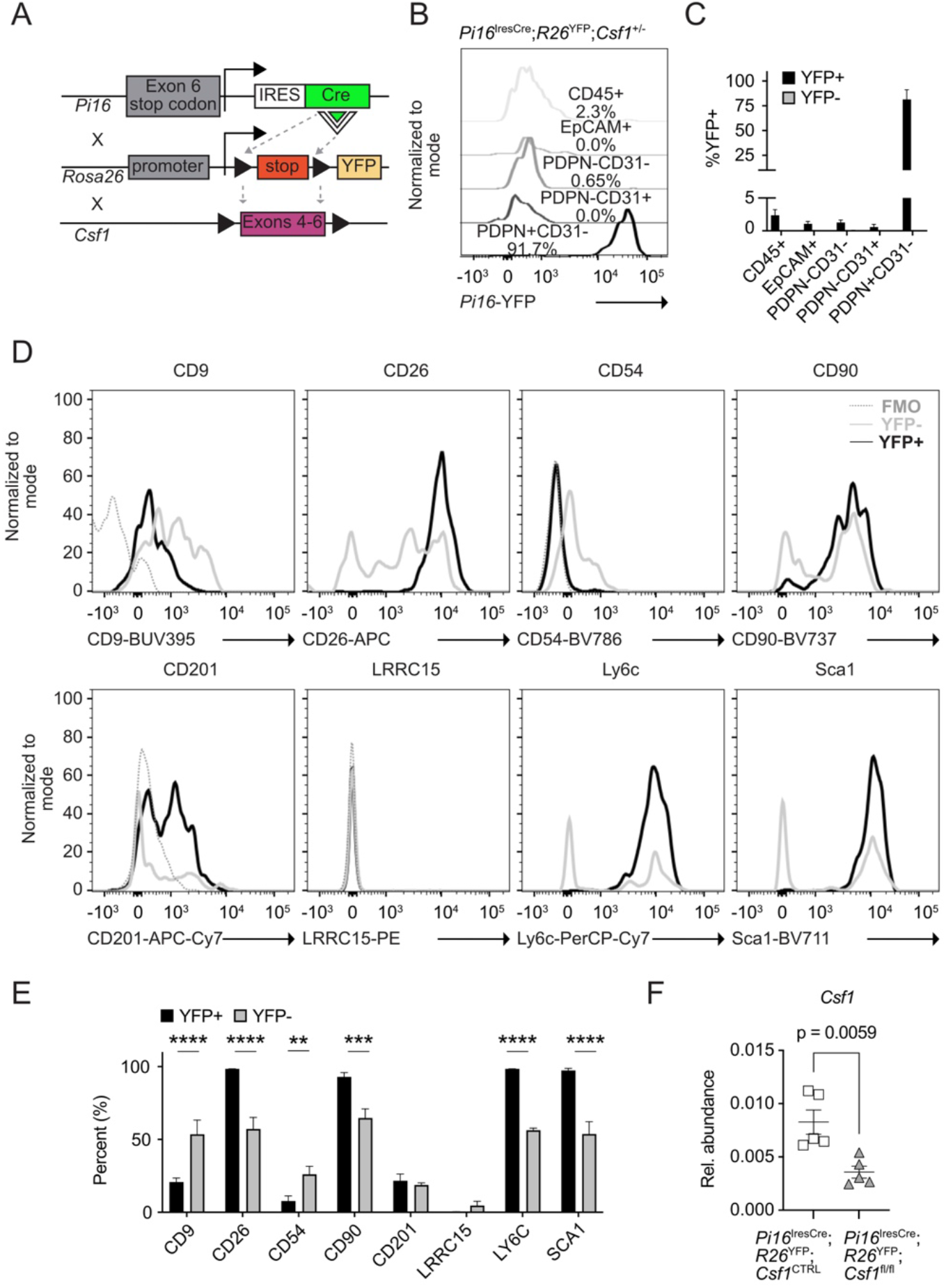
Pi16^+^ fibroblasts are the dominant substate of skin fibroblasts and produce Csf1. **(A)** graphical schematic of *Pi16*^*IresCre*^*;R26*^*YFP*^*;Csf1*^*fl/fl*^ mouse. **(B)** representative gating and **(C)** frequency of YFP^+^ (n=7) and YFP^-^ cell (n=3) populations in the skin of *Pi16*^*IresCre*^*;R26*^*YFP*^*;Csf1*^*+/-*^ mice. **(D)** representative gating of CD9, CD26, CD54, CD90, CD201, LRRC15, LY6C, SCA1 abundance on PDPN^+^YFP^+^ and PDPN^+^YFP^-^ fibroblasts in the skin of *Pi16*^*IresCre*^*;R26*^*YFP-CTRL*^ mice (n=3). **(E)** Quantification of percent CD9, CD26, CD54, CD90, CD201, LRRC15, LY6C, and SCA1 in the skin of *Pi16*^*IresCre*^*;R26*^*YFP-CTRL*^ *mice* (n=3). **(F)** qPCR analysis of *Csf1* in the skin of *Pi16*^*IresCre*^;*R26*^YFP^;*Csf1*^fl/fl^ (n=5) and *Pi16*^*IresCre*^;*R26*^YFP-CTRL^ (n=5) mice **(A-D)**, Cells were gated on live. **(E)** Statistics were calculated using Two-way ANOVA. **(F)** Statistics were calculated using an unpaired, two-tailed Student’s t-test. **p<0.01, ***p<0.001, ****p<0.0001. Data are mean ± s.e.m.

### Pi16^+^ fibroblasts maintain a skin macrophage niche

We hypothesized that disruptions in *Pi16*^+^ fibroblast-derived *Csf1* signaling would impact macrophage homeostasis in the skin. Previous single-cell multi-omic profiling demonstrated that skin macrophages can be broadly grouped as CD64^+^ or CD11c^+^, with increased expression of tissue-resident macrophage genes (i.e., *Timd4*/TIM4, *Mrc1*/CD206)^21^ or monocyte-associated genes (i.e. *Ccr2*/CCR2), respectively^18^. We subsequently measured changes in F4/80^+^, CD64^+^, and CD11c^+^ skin macrophage populations in *Pi16*^IresCre^;*R26*^YFP^;*Csf1*^fl/fl^ and control mice by flow cytometry. A significant decrease in the frequency (1.3-fold) and number (4.3-fold) of total F4/80^+^ macrophages was observed in *Pi16*^IresCr*e*^;*R26*^YFP^;*Csf1*^fl/fl^ mice compared to *Pi16*^IresCre^;*R26*^YFP^;*Csf1*^CTRL^ mice (**Fig. 2A-B)**. Significant decreases in both CD64^+^ (7.9-fold) and CD11c^+^ (3.4-fold) macrophages per milligram of tissue were also observed in the skin *Pi16*^IresCre^;*R26*^YFP^;*Csf1*^fl/fl^ mice compared to *Pi16*^IresCre^;*R26*^YFP^;*Csf1*^CTRL^ animals (**Fig. 2A,C)**. No changes were observed in Ly6c^+^Cd11b^+^ bone marrow monocytes in *Pi16*;*R26*^YFP^;*Csf1*^fl/fl^ mice compared to control animals (**Supplemental Fig. 1A-B**). These results demonstrate that *Pi16*^+^ fibroblast-derived *Csf1* is critical, local source for maintaining skin macrophages.

**Figure 2.**
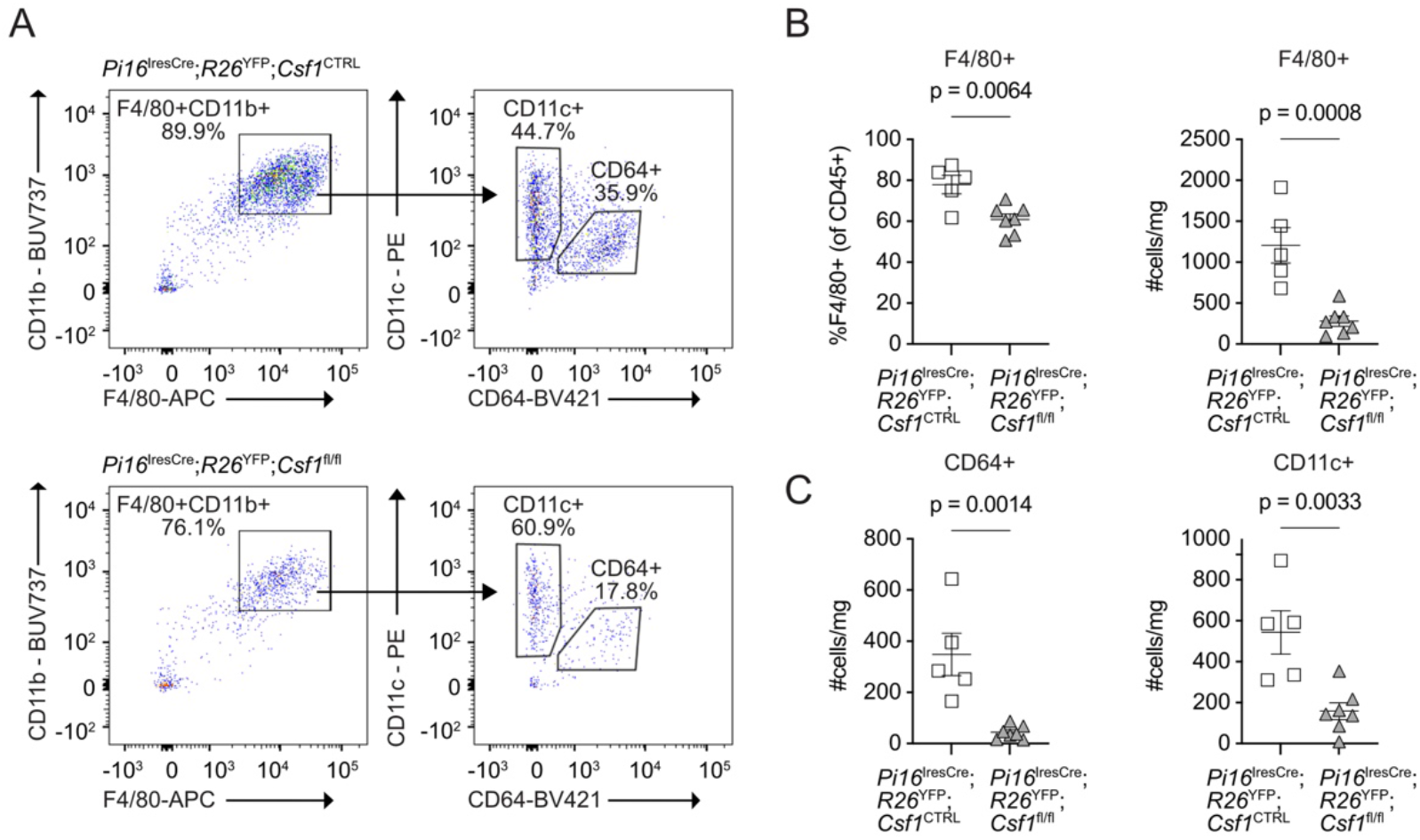
Pi16^+^ fibroblasts maintain a skin macrophage niche. **(A)** representative gating of CD45^+^Gr1^-^F4/80^+^CD11b^+^CD64^+^CD11c^+^ macrophages in the skin. **(B)** frequency and enumeration of CD11b^+^F4/80^+^ macrophages and **(C)** CD64^+^ and CD11c^+^ macrophages in the skin of *Pi16*^*IresCre*^*;R26*^*YFP-CTRL*^ (n=6) and *Pi16*^*IresCre*^*;R26*^*YFP*^*;Csf1*^*fl/fl*^ (n=8) mice. Cells were gated on live, CD45^+^, Gr1^-^, F4/80^+^. Statistics were calculated using an unpaired, two-tailed, Student’s t-test. Data are mean ± s.e.m.

### Loss of skin macrophages drives PDPN^+^YFP^+^ fibroblast expansion in the skin

Our previous work demonstrated that the progressive loss of CD64^+^ and CD11c^+^ macrophage populations altered fibroblast transcriptional states and population dynamics in the skin^18^. Therefore, we hypothesized that constitutive loss of *Csf1* in *Pi16*^+^ fibroblasts, in addition to eliciting diminishment of both skin macrophage populations, would increase the abundance of skin fibroblasts. Flow cytometric analysis demonstrated an increased frequency (2.1-fold) and number of PDPN^+^ fibroblasts per milligram (1.5-fold) in the skin of *Pi16*^IresCre^;*R26*^YFP^;*Csf1*^fl/fl^ mice relative to control animals (**Fig. 3A-B)**. PDPN^+^YFP^+^ fibroblast frequency (1.2-fold) and number per milligram (2.1-fold) were also increased *Pi16*^IresCre^;*R26*^YFP^;*Csf1*^fl/fl^ compared to control animals (**Fig. 3A,C**). The number of PDPN^+^YFP^+^ CD26^+^ (3.5-fold), LY6C^+^ (3.4-fold), and SCA1^+^ (3.5-fold) fibroblasts were significantly increased (**Fig. 3D)**. No changes the percent of PDPN^+^YFP^+^ cells expressing CD26, LY6C, or SCA1 cells were observed between *Pi16*^IresCre^;*R26*^YFP^;*Csf1*^fl/fl^ and control animals but loss of *Csf1* coincided with a diminishment of CD90 expression (**Fig. 3E)**. These results suggest that CD64^+^ and CD11c^+^ macrophage populations are key arbiters of *Pi16*^+^ population dynamics in the skin and minimally affect the phenotype of these cells.

**Figure 3.**
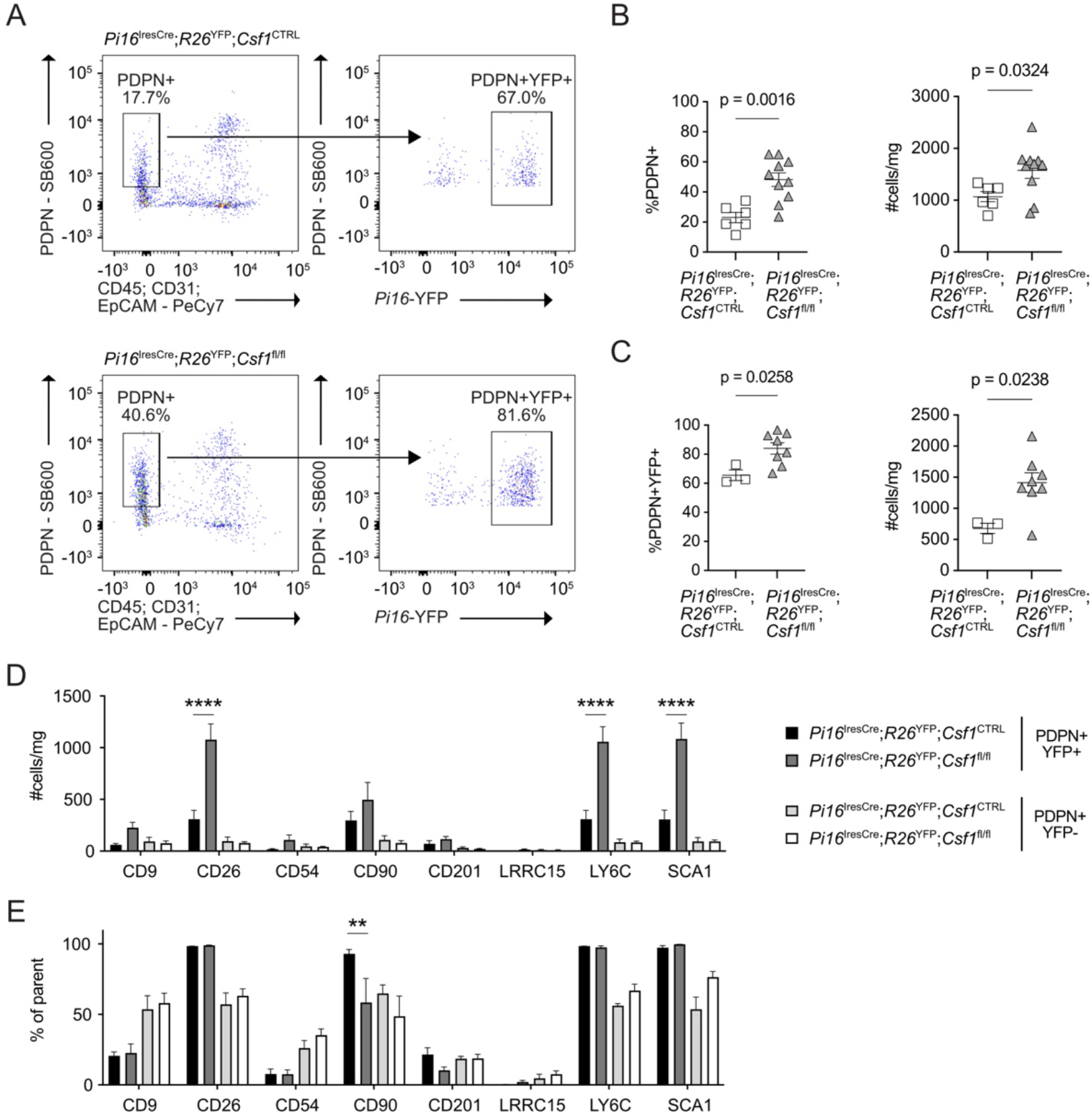
Conditional deletion of Csf1 in Pi16^+^ fibroblasts results in PDPN^+^YFP^+^ fibroblast expansion in the skin. **(A)** representative gating of CD45^-^CD31^-^EpCAM^-^PDPN^+^YFP^+^ fibroblasts in the skin of *Pi16*^*IresCre*^*;R26*^*YFP-CTRL*^ and *Pi16*^*IresCre*^*;R26*^*YFP*^*;Csf1*^*fl/fl*^ mice. **(B)** frequency and enumeration of PDPN^+^ fibroblasts in the skin of *Pi16*^*IresCre*^*;R26*^*YFP-CTRL*^ (n=6) and *Pi16*^*IresCre*^*;R26*^*YFP*^*;Csf1*^*fl/fl*^ (n=10) mice. **(C)** enumeration and frequency of PDPN^+^YFP^+^ fibroblasts in the skin of *Pi16*^*IresCre*^*;R26*^*YFP-CTRL*^ (n=3) and *Pi16*^*IresCre*^*;R26*^*YFP*^*;Csf1*^*fl/fl*^ (n=8) mice. **(D)** Enumeration of the number and **(E)** frequency of parent of PDPN^+^YFP^+^ and PDPN^+^YFP^-^CD9, CD26, CD54, CD90, CD201, LRRC15, LY6C, and SCA1 in the skin of *Pi16*^*IresCre*^*;R26*^*YFP-CTRL*^ (n=3) and *Pi16*^*IresCre*^*;R26*^*YFP*^*;Csf1*^*fl/fl*^ (n=8) mice. (B, C) Statistics were calculated using an unpaired, two-tailed, Student’s t-test. (D) Statistics were calculated using Two-way ANOVA. **p<0.01, ****p<0.0001. Data are mean ± s.e.m.

### Altered fibroblast:macrophage circuits dysregulate skin architecture and wound healing

Based on our previous work, we predicted that the expansion of PDPN^+^YFP^+^ skin fibroblasts due to CD64^+^ and CD11c^+^ macrophage loss would also be coincident with increased skin thickness^18^. Therefore, we performed histological assessment of Hematoxylin & Eosin (H&E)-stained flank skin tissue sections. We observed increases in the dermis (1.3-fold), panniculus muscle (PM; 1.5-fold), and fascia (1.7-fold) compartments of *Pi16*^IresCre^;*R26*^YFP^;*Csf1*^fl/fl^ relative to control mice (**Fig. 4A-B**). Immunohistochemistry analysis of skin tissue sections demonstrated increased fibroblast frequency (1.8-fold) in the fascia (**Fig. 4C-D**), but not in epidermis or dermal white adipose tissue (dWAT). Loss of F4/80^+^ macrophages (2.0-fold) per square micrometer (mm^2^) of fascia was also observed *Pi16*^IresCre^;*R26*^YFP^;*Csf1*^fl/fl^ compared to control animals (**Fig. 4e)**.

**Figure 4.**
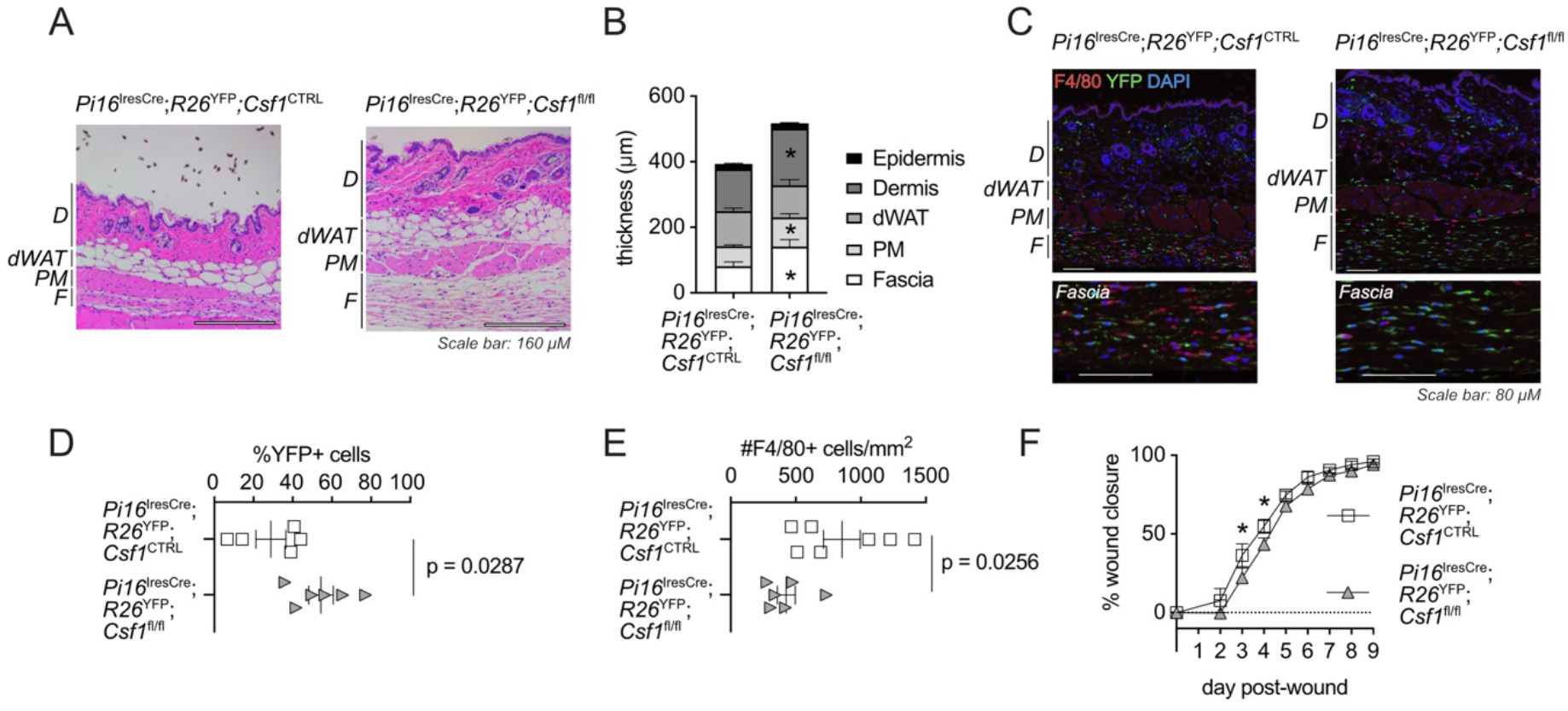
Dysregulated skin architecture and wound healing kinetics in Pi16^IresCre^;R26^YFP^;Csf1^fl/fl^ mice. **(A)** representative images and **(B)** quantification of skin tissue compartment thickness in hematoxylin and eosin-stained histological sections of *Pi16*^*IresCre*^*;R26*^*YFP-*CTRL^ (n=8) and *Pi16*^*IresCre*^*;R26*^*YFP*^*;Csf1*^*fl/fl*^ (n=6) mice. **(C)** representative IHC images of F4/80^*+*^ macrophages (red), YFP^+^ fibroblasts (green), and DAPI nuclei (blue) in the skin *of Pi16*^*IresCre*^*;R26*^*YFP-CTRL*^ and *Pi16*^*IresCre*^*;R26*^*YFP*^*;Csf1*^*fl/fl*^ mice. **(D)** frequency of YFP+ fibroblasts in the skin of *Pi16*^*IresCre*^*;R26*^*YFP-CTRL*^ (n=5) and *Pi16*^*IresCre*^*;R26*^*YFP*^*;Csf1*^*fl/fl*^ (n=6) mice. **(E)** enumeration of the number of F4/80^*+*^ macrophages per mm^2^ of skin in *Pi16*^*IresCre*^*;R26*^*YFP-CTRL*^ (n=7) and *Pi16*^*IresCre*^*;R26*^*YFP*^*;Csf1*^*fl/fl*^ (n=6) mice. **(F)** percent closure at days 0-9 post wound healing in of *Pi16*^*IresCre*^*;R26*^*YFP-*CTRL^ (n=6) and *Pi16*^*IresCre*^*;R26*^*YFP*^*;Csf1*^*fl/fl*^ (n=7) mice. (**B**,**D**,**E)**, Statistics were calculated using an unpaired, two-tailed, Student’s t-test. **(G)** Statistics were calculated using unpaired Two-way ANOVA. Data are mean ± s.e.m.

Based on these observations in the steady state, our previous work demonstrating that *Dpt*^+^ fibroblasts promote efficient wound healing^18^ and a report demonstrating that skin fascia fibroblast progenitors are essential for effective wound healing^22^, we hypothesized that the *Pi16*^+^ fibroblast-macrophage axis may be critical in the wound healing process. We examined wound healing kinetics in *Pi16*^IresCre^;*R26*^YFP^;*Csf1*^fl/fl^ compared to control animals using an unsplinted model. We observed a significant decrease in percent closure at day 3 (average percent closure: control, 36.4%; and *Pi16*^IresCre^;*R26*^YFP^;*Csf1*^fl/fl^, 23.3%) and 4 (average percent closure: control, 55.0%; and *Pi16* ^IresCre^;*R26*^YFP^;*Csf1*^fl/fl^, 45%) post wound in *Pi16*^IresCre^;*R26*^YFP^;*Csf1*^fl/fl^ compared to control mice (**Fig. 4F-H)**, suggesting that the *Pi16*^+^ fibroblast-macrophage axis accelerates early wound healing. These findings suggest that macrophage loss can elicit changes in the population dynamics of fibroblasts to modify tissue architecture in the presence and absence of inflammation.

### Pi16^+^ fibroblast frequency is suppressed in the skin during systemic sclerosis

Systemic scleroderma (SSc) is characterized by skin fibrosis, thickening, and fibroblast to myofibroblast conversion^23^. In addition to demonstrating that loss of *Pi16*^+^ fibroblasts resulted in dermal and fascial thickening (**Fig. 4C-D**), we previously demonstrated that dysregulation of the *Dpt*^+^ fibroblast-macrophage axis was associated with SSc pathogenesis^18^. Therefore, we hypothesized that the *Pi16*^+^ fibroblast:macrophage circuit is critical for maintaining human skin homeostasis. To delineate the role of the *Pi16*^+^ fibroblast substate in skin health and disease, we examined a publicly available dataset of skin samples from healthy controls (n=56) and patients with limited (n=38) or diffuse SSc (n=48), where *CSF1* expression is positively associated with disease pathogenesis^18,20^. Our analysis of skin fibroblast populations demonstrated that *PI16* expression was significantly enriched (*logFc>2, adj p-val<0*.*01*) in Scleroderma-Associated Fibroblasts (ScAF)^20^. The *PI16+*/ScAF cluster differentially expressed *LGR5, DPP4, WISP2*, and *SLPI*, among other genes (**Fig. 5A and Supplementary Table 1**). We observed a decrease in the *PI16*^+^/ScAF cluster in patients with lSSc and dSSc compared to healthy controls and a diminishment of the *PI16+*/scAF cellular neighborhood as determined by Milo analysis^24^, consistent with previous reports that note a diminishment in scAF with SSc progression^20^ (**Fig. 5B-C**). These results suggest that *PI16*^*+*^ fibroblasts are a component of the ScAF phenotype and that loss of *PI16+* fibroblasts is associated with SSc disease progression.

**Figure 5.**
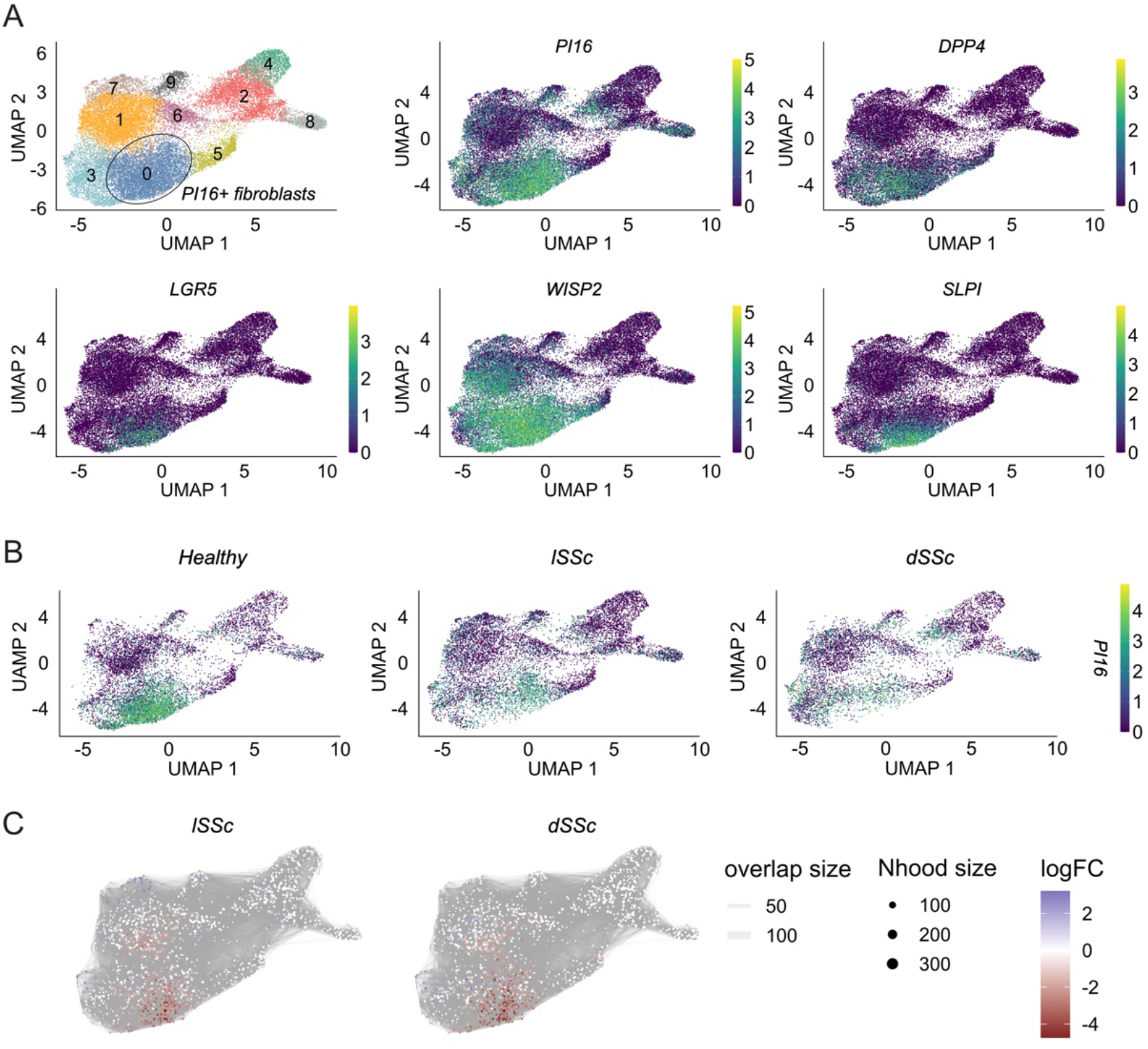
PI16^+^ fibroblast frequency in healthy and SSc human skin. **(A)** UMAP visualization of single cells (top left) and dot plots for *PI16, DPP4, LGR5, WISP2*, and *SLPI* from healthy controls (n=56) and patients with lSSc (n=38) or dSSc (n=48) in human skin tissue, reanalyzed from a previous study^20^. *PI16*^+^ fibroblasts are annotated. **(B)** *PI16* expression across skin cell clusters in healthy controls (n=56) and patients with lSSc (n=38) or dSSc (n=48) in human skin tissue. **(C)** Milo differential abundance analyses of *PI16*^+^ fibroblasts healthy controls (n=56) and patients with lSSc (n=38) or dSSc (n=48) in human skin tissue.

## Discussion

Fibroblasts and macrophages are found across all tissues and have been hypothesized to be interacting partners for a century^10,25^. We recently revealed that *Dpt*^+^ fibroblasts and macrophage exist in a push-pull circuit that is essential for skin integrity^18,26^. Despite this, whether *Pi16*^+^ fibroblasts comprise a macrophage niche, as well as their impact on tissue architecture and function was not well understood. In this study, we provide corroborating evidence of reciprocal regulation between fibroblasts and macrophages in the skin and demonstrate that *Pi16*^+^ skin fibroblasts and macrophages exist in a two-cell circuit anchored by fibroblast-derived *Csf1* that is essential for steady-state tissue architecture and wound healing.

We performed flow cytometric analyses and demonstrated that *Pi16*^+^ fibroblasts represent ∼80% of fibroblasts in the skin. These fibroblasts had high abundance of markers *Dpp4*/CD26, *Thy1*/CD90, *Ly6c1*/Ly6c, and *Ly6a*/Sca1, similar to previously described reticular fibroblasts (*PI16*^+^*CD34*^+^*DPP4*^*+*^)^27^; adventitial stromal cells (ASC; *Pi16*^+^*Ly6a*^+^)^28^; fibroinflammatory progenitor cells (FIP; *Pi16*^+^*Dpp4*^+^)^29^; T Helper 2 (TH2)-interacting fascial fibroblasts (TIFFs; *IL13ra1*^+^*Pi16*^+^*Anxa3*^+^)^30^; and (*Dpt*^+^*Pi16*^+^*Dpp4*^+^) *Engrailed-1* (*En1*)^31^ cells^32^. *En1*+ fibroblasts, human reticular fibroblasts, and TIFF are found in the fascia of the skin and regulate tissue architecture and wound healing^27,30,31^. *En1*^+^ fibroblasts have been previously shown to be the dominant cell type in adult skin and produce collagen to influence architecture and wound repair^31^. Skin reticular fibroblasts and express members of the Kruppel like factor (KLF) transcription factor family^27^ which represses myofibroblast- and inflammation-associated transcription profiles^33^. TIFFs expand in response to TH2-derived cytokines to form subcutaneous fibrous bands^30^, highlighting their immune-facing role. In the lungs, ASC dictate the localization of type 2 innate lymphoid cells (ILC2) in the steady-state and allergic asthma^28^. FIP express high levels of monocyte chemokines *Ccl2, Cxcl10*, and *Cxcl2*^29^ in adipose tissue. Collectively, these reports suggest that the *Pi16*^+^ fibroblast armamentarium utilizes both regenerative and immunomodulatory mechanisms for unique tissue functions in steady-state and disease.

We demonstrate that constitutive deletion of *Csf1 in Pi16*^*+*^ fibroblasts resulted in the loss of tissue *Csf1*, as well as the CD64^+^ and CD11c^+^ macrophage niche, representative of tissue-resident and monocyte-derived populations, respectively^18^. We observed concurrent expansion of the number of PDPN^+^ and (*Pi16*^+^) PDPN^+^YFP^+^ fibroblasts per milligram of tissue with only a minimal shift in fibroblast surface marker expression, suggesting that macrophages restrain fibroblast expansion but minimally shape *Pi16*^+^ fibroblast phenotype in the skin. Orthogonal analyses in skin fascia by immunofluorescence and histological demonstrated that F4/80^+^ cells were decreased, whereas YFP^+^ cells and tissue thickness were increased, consistent with previous reports^27,30,31^. Expansion may be driven by dysregulated spatiotemporal Hippo-YAP signaling, as previously described in mice with a conditional deletion of *Csf1* in *Dpt*^+^ fibroblasts^18^ or skin edema resulting from CSF1R blockade^34^. Nonetheless, our results demonstrate that fibroblast:macrophage circuits are critical for the collectively topography of the skin, even in the absence of inflammation. During injury, we demonstrated that deletion of *Csf1* in *Pi16*^+^ fibroblasts resulted in a modest, but significant delay in early wound healing. These findings are aligned with our previous study investigating skin wounding dynamics following *Csf1* deletion in *Dpt*^+^ fibroblasts^18^ and those which deplete skin *Mrc1*^*+*^ macrophages^6,8^ indicating that the fibroblast:macrophage circuit is critical for tissue regeneration as well as homeostasis.

It has been demonstrated that organs subject to cycles of mechanical compression and distension, including the skin, lungs, heart, gastrointestinal tract, and musculoskeletal systems are surrounded by a supportive collagen and fluid filled sinus, often termed the reticular interstitium, as opposed to discrete tissue-specific nomenclature for microanatomical compartments, i.e., skin fascia/adventitia^35^. As *Pi16*^+^ fibroblasts are found across tissues^17^, additional studies are required to determine the roles and localization of *Pi16*^+^ fibroblasts in other organs, as well as how their interactions with other tissue-resident immune cells, including macrophages and long-lived lymphocytes, shape architecture and function in the steady-state.

The precise role(s) of *Pi16*^+^ fibroblasts are still evolving and being uncovered. Here, we reveal a key role for *Pi16*^+^ fibroblast-derived *Csf1* in shaping the macrophage setpoint and integrity of the skin. A precursor role for these cells during inflammation in the skin has been suggested^27,33^. Other work suggests that these cells may have a role in shaping human immune responses in secondary lymphoid organs^36^ and may be a positive prognostic indicator across cancer indications^37^. Our skin-focused analysis of *PI16*^+^ fibroblasts suggests these cells are inversely correlated with disease severity in SSc. The mechanism here may owe to the ability of these cells to serve as a precursor cell to more activated fibroblasts. A limitation of this approach is the use of *PI16* alone, which does not account for alterations within the *Pi16*^+^ fibroblast cell state that may occur absent differentiation, which has been noted in other contexts^38^. Nonetheless, preserving the native *Pi16*^+^ cell state through transcription factor engagement, such as enforcing expression of KLF4, has been recently proposed to harness the therapeutic potential of *PI16*^+^ fibroblasts^27,33^. Future approaches like this may be a valid means to utilize *PI16*^+^ fibroblasts to treat diseases including SSc, and possibly others.

## Supplemental figures

**Supplemental figure 1.**
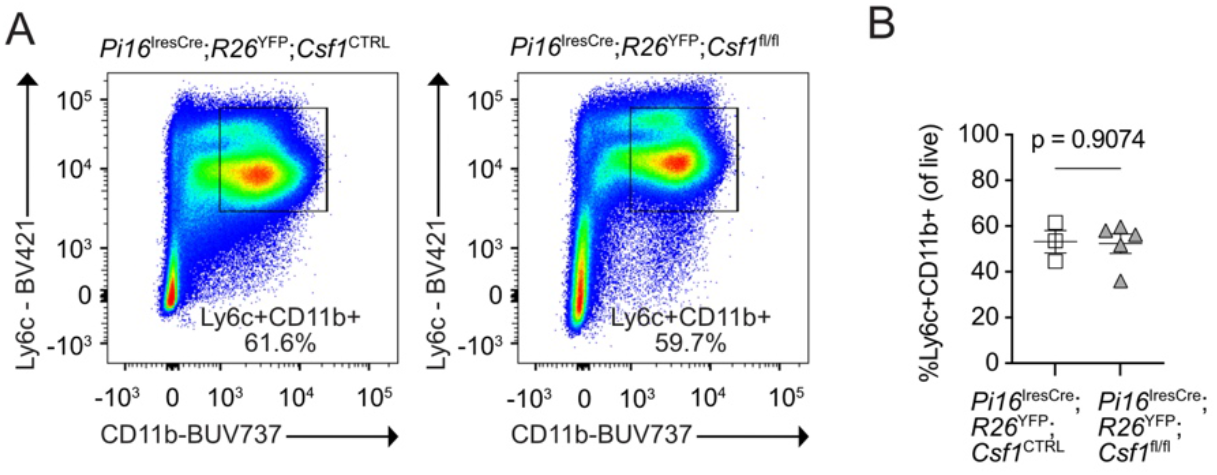
Constitutive deletion of Csf1 in Pi16^+^ fibroblasts does not alter bone marrow macrophage homeostasis. (**A)** representative gating and **(B)** frequency of CD45^+^Ly6c^+^CD11b^+^ monocytes in the bone marrow of *Pi16*^*IresCre*^*;R26*^*YFP-CTRL*^ (n=3) and *Pi16*^*IresCre*^*;R26*^*YFP*^*;Csf1*^*fl/fl*^ (n=5) mice. Statistics were calculated using an unpaired, two-tailed, Student’s t-test. Data are mean ± s.e.m.

**Supplemental figure 2.**
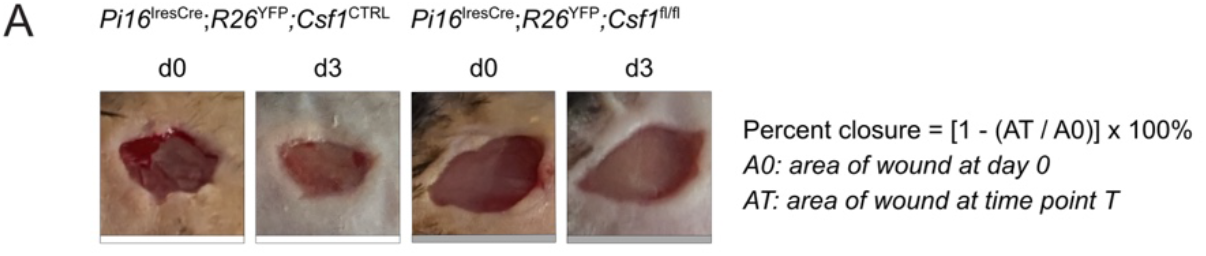
Representative images of an unsplinted mouse model of wound healing. (**A)** representative images at day 3 post-wound and percent change calculations for an unsplinted model of full thickness wound healing.

## Acknowledgments

We are grateful to the staff at TCP for animal husbandry and colony maintenance assistance. We thank the Centre for Immune Analytics (RRID:SCR_027612) in the Department of Immunology at the University of Toronto for assistance with flow cytometry. We thank the Advanced Optical Microscopy Facility at University Health Network.

## Supplementary material

Supplemental material is available online.

## Data availability

The data underlying this article will be shared on reasonable request to the corresponding authors.

